# Fatty Acid Vesicles as Hard UV-C Shields for Early Life

**DOI:** 10.1101/2023.01.01.522439

**Authors:** Iván Lechuga, Karo Michaelian

## Abstract

Theories on life’s origin generally acknowledge the advantage of a semi-permeable vesicle (protocell) for enhancing the chemical reaction-diffusion processes involved in abiogenesis. However, more and more evidence indicates that the origin of life concerned the photo-chemical dissipative structuring of the fundamental molecules under UV-C light. In this paper, we analyze the Mie UV scattering properties of such a vesicle made from long chain fatty acids. We find that the vesicle could have provided early life with a shield from the faint, but dangerous, hard UV-C ionizing light (180-210 nm) that probably bathed Earth’s surface from before the origin of life and until perhaps 1,500 million years after, until the formation of a protective ozone layer as a result of the evolution of oxygenic photosynthesis.

## 1 Introduction

The physical and chemical properties of vesicles made up of amphiphilic molecules such as fatty acids, isoprenes, and phospholipids have been analyzed in detail in relation to their utility as micro-environments for enhancing reaction-diffusion processes occurring in prebiotic cells at the origin of life [1, 2, 3, 4]. Amphiphilic membranes play a role in energy transduction, ion transport, and concentrating molecules, thereby increasing reaction rates [2]. The membrane also allows regulated exchange of incoming and outgoing material [5, 6] leading to the formation of gradients, which can be useful as energy sources themselves [7].

However, more and more evidence indicates that the origin of life concerned not only chemical reactions, but also photochemical reactions. In particular, it has been hypothesized that the fundamental molecules of life were formed through dissipative structuring from precursors under the surface UV-C light environment of the Archean [5, 8, 9, 10, 11, 12]. Assuming such a scenario, it would be important to understand how the optical properties of the vesicle could have affected the UV-C photochemical dissipative structuring of the fundamental molecules within. This paper presents a first (as far as the authors are aware) analysis of the optical properties of such vesicles at UV-C wavelengths and how these properties could have influenced the molecular dissipative structuring processes.

The dissipative structuring of material under an externally imposed generalized thermodynamic potential is a well-established non-equilibrium thermodynamic result which leads to the complexation of physical-chemical systems in such a manner as to increase the dissipation of the imposed potential [13]. The Thermodynamic Dissipation Theory for the Origin and Evolution of Life [5, 8, 9, 10, 12, 14, 15, 16] identifies this generalized thermodynamic potential for life’s dissipative structuring involved in the origin of life as the solar photon potential of UV-C wavelengths between 205 and 285 nm. In this region, photons can break and remake covalent bonds in carbon-based molecules but rarely cause molecular dissociation through ionization. This photon potential was available at Earth’s surface before the origin of life (~ 3.9 Ga) and persisted for at least 1200 million years until the emergence of oxygenic photosynthesis and an ensuing ozone layer at about 2.7 Ga.

The thermodynamic dissipation theory for the origin of life proposes that the fundamental molecules of life (those in the three domains of life - nucleic acids, amino acids, fatty acids, sugars, cofactors, etc.) were initially UV-C pigments [10] dissipatively structured on the ocean surface from common precursors such as HCN, cyanogen, CO_2_, and water under the UV-C photon potential [5, 12, 15, 17]. It is assumed that prebiotic vesicles would form from these dissipatively structured fatty acids and isoprenes at the ocean surface, through surface agitation and natural Gibb’s free energy minimization.

In this paper, we investigate the optical properties of pre-biotic vesicles made from fatty acids given their physical-chemical properties which would have been required for vesicle stability under the ambient conditions of Earth’s Archean ocean surface. Specifically, we determine the vesicle wavelength-dependent absorption and scattering given the requirements of stability against high temperature, pH variation, and salt flocculation. In particular, we analyze the possibility that vesicles could have acted as a Mie scattering particles for the high energy ionizing UV-C photons in the 180-210 nm region (labeled here as “hard” UV-C photons), thereby preventing damage to the fundamental molecules of life, which we assume were dissipatively structured within the vesicle with basically “soft” UV-C photons (defined here as having wavelengths between 245 and 275 nm). We find that an important photo-protective effect against hard UV-C photons indeed occurs for specific vesicle radii which, interestingly, are concordant with the size distribution of oldest bacteria found in the Archean fossil record at 3.49, 3.45 and 3.2 Ga [18].

Furthermore, we suggest that, as a result of UV-C light back-scattering from vesicles, followed by total internal reflection at the ocean surface, vesicles could thereby increase the amount of circularly polarized UV-C light available to the molecules confined within neighboring vesicles. This has relevance to the origin of life since we have proposed elsewhere [14, 16] that the homochirality of life arose from the DNA or RNA UV-C photon-induced denaturing resulting from the dissipation of these soft UV-C photons in double strand DNA and RNA, in conjunction with the asymmetry in this light-induced denaturing due to the small morning/afternoon temperature difference of the ocean surface.

## 2 Physical, Chemical, and Optical Properties of Archean Vesicles

### 2.1 Properties of Fatty Acid Vesicles at the Origin of Life

Natural fatty acids are formed from a carboxyl head group with a hydrocarbon tail of from 4 to 40 carbon atoms [19]. Empirical evidence indicates that the earliest ancestors of the modern cellular membranes were devoid of the relatively more complex phospholipids [20, 21]. Furthermore, simple fatty acids are easily produced as small chains of ethylene (C_2_H_4_) through activated Fischer-Tropsch, or soft UV-C photochemical, polymerization [15]. Ethylene itself can be produced from the reduction of CO_2_ or CO in water, or from UV photolysis of methane [15]. Today, fatty acids are found in the membranes of the cell walls of organisms from all three domains of life [21].

From the perspective of the thermodynamic dissipation theory for the origin of life, we have suggested that conjugated fatty acids of about 18 carbon atoms were prominent among the fatty acids which formed the first prebiotic vesicles [15]. Vesicles of fatty acids of these lengths are stable at the high temperature of early Archean surface (~ 85°C [22, 23]). The critical vesicle concentration (CVC) of these fatty acids required for vesicle formation decreases as the alkyl chain length increases, since longer chains favor increased packing of bilayers. Triple conjugated forms of these fatty acids absorb strongly near 260 nm, which is at the peak of the UV-C surface photon spectrum available throughout the Archean (Fig. 1).

**Figure 1.**
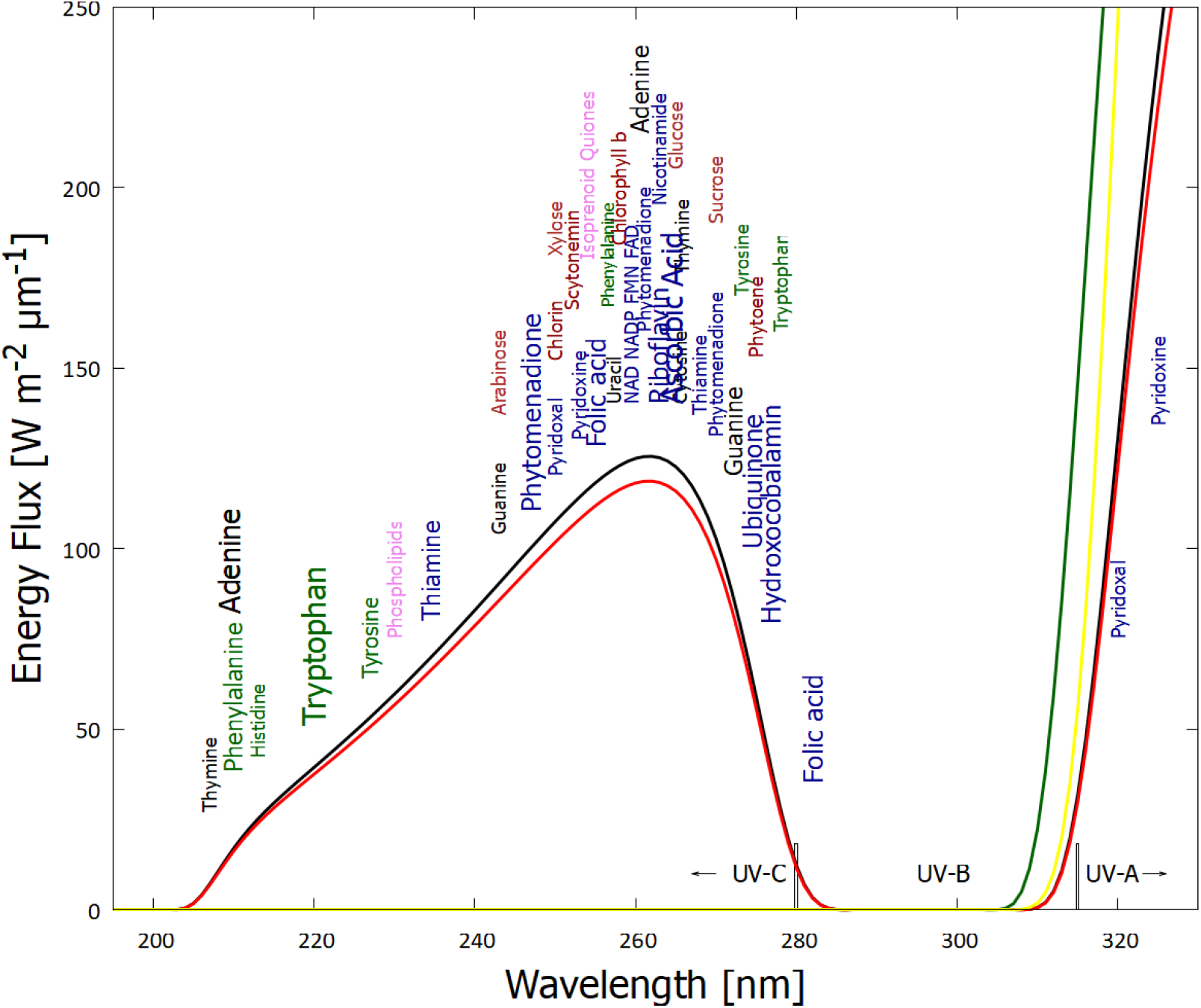
The spectrum of UV light available at Earth’s surface before the origin of life at approximately 3.9 Ga and until at least 2.9 Ga (curves black and red respectively) (perhaps even extending throughout the entire Archean until 2.5 Ga [24]). Atmospheric CO_2_ and perhaps some volcanic H_2_S were responsible for absorption at wavelengths shorter than ~205 nm, and atmospheric aldehydes (common photochemical products of CO_2_ and water) absorbed between about 285 and 310 nm [25], approximately corresponding to the UV-B region. Around 2.2 Ga (green curve), UV-C light at Earth’s surface was completely extinguished by oxygen and ozone resulting from organisms performing oxygenic photosynthesis. The yellow curve corresponds to the present-day surface spectrum. Energy fluxes are for the sun at the zenith. The names of the fundamental molecules of life are plotted at their wavelengths of maximum absorption; nucleic acids (black), amino acids (green), fatty acids (violet), sugars (brown), vitamins, co-enzymes and cofactors (blue), and pigments (red) (the font size roughly corresponds to the relative size of their molar extinction coefficient). Adapted from [10].

Indeed, evidence in the Archean fossil record, and also in modern oceans, shows a predominance of fatty acids with an even number of carbon atoms, particularly a predominance of 16 and 18 carbons [26, 27]. The ubiquity of chains with an even number of carbon atoms could be explained if the early Archean fatty acids were formed through the polymerization of ethylene (C_2_H_4_). In fact, there is a very simple photochemical dissipative structuring route involving the polymerization of ethylene to form fatty acids, starting from CO or CO_2_ saturated water [15, 28].

The melting temperature of vesicles generally increase with fatty acid carbon tail length and the degree of hydrogen saturation, with the greater influence being due to saturation. Long chain fatty acids which are partially unsaturated (and conjugated) could form stable vesicles at the high temperatures of the early Archean. For example, fully conjugated (triple) linolenic acid (cis-9,cis-12,cis-15 18:3) has a strong and wide absorption band at 269 nm and a vesicle melting temperature of around 85 ° C (Earth’s average surface temperature as obtained from isotopic evidence in sediments dating back to the origin of life [22, 23]).

### 2.2 Photochemical Dissipative Structuring of Fatty Acids

We have suggested that the formation of fatty acids was most likely through soft UV-C photochemical dissipative structuring in CO_2_ saturated water at the ocean surface, leading to ethylene and its polymerization [12, 15]. In support of this suggestion is the fact that the polymerization rates of ethylene are two orders of magnitude greater at 254 nm (UV-C) than at 365 nm (UV-A) [29].

Another possible photo-induced route to fatty acids synthesis in the UV-C and UV-B regions starts with formaldehyde under this light and ZnO and TiO_2_ acting as photocatalysts, leading to the formation of 2 to 5 carbon fatty acids, as well as to other fundamental molecules of life [30]. For example, the route to fatty acid formation includes the oligomerization of HCN into diaminomaleonitrile DAMN which requires absorption in the UV-C region between 205 and 285 nm, similar to the photochemical dissipative structuring routes of the purines proposed elsewhere by the authors [5, 6].

It is known that saturated (non-conjugated) fatty acids absorb very little in the UV-C region, except for dissociation below ~ 180 nm and a small absorption peak from the carboxyl head group at 207 nm [31]. UV-C light between 205 and 285 nm can, however, induce de-protonation of fatty acids leading to conjugation of carbon bonds in the acyl tail [32]. After several deprotonation events and migration of the double bonds, the fatty acids arrive at a conjugated configuration; diene for two double bonds with absorption at 233 nm, triene for three double bonds with absorption at 269 nm, and tetraene for those with four double bonds giving absorption at 310-340 nm. All of these wavelengths lie within the likely Earth’s Archean surface photon spectrum (Fig. 1). Furthermore, conjugated fatty acids have conical intersections [33] allowing rapid (sub-picosecond) dissipation of the photon-induced excited state energy into harmless heat, which is yet another strong indication of their production at the origin of life through UV-C dissipative structuring [5, 12].

The same UV-C photons can also induce cross-linking between adjacent tails of fatty acids at the site of double bounds, resulting in a reduction in the average conjugation number [34]. A stationary state distribution of the fatty acid conjugation number, with a maximum at triple conjugation (giving absorption at 269 nm, near the peak in the UV-C surface spectrum - Fig. 1) would thus arise under the constant UV-C flux [15].

The steps involved in the dissipative structuring of fatty acids can thus be summarized as follows [15]; i) UV-C-induced reduction of CO_2_ or CO in water saturated with these, forming ethylene, ii) UV-C-induced polymerization of ethylene to form long hydrocarbon tails of an even number of carbon atoms (e.g. 18C), iii) oxidation and hydrolysis events to stop the growth of the chain and form the carboxyl head group, respectively, iv) UV-C induced deprotonation of the tails to form double carbon-carbon bonds, v) double bond migration to give a conjugated diene, triene, or tetraene molecule with a conical intersection. The conical intersection provides sub-picosecond decay of electronic excited state energy into harmless heat, preventing further photochemical reactions and, thus, providing photochemical stability, and, most importantly, from our perspective of dissipative structuring, photon dissipative efficacy.

### 2.3 Vesicle formation

The problems associated with fatty acids vesicle formation and stability are; 1) the narrow range of alkaline pH in which vesicles spontaneously form, 2) salt flocculation (agglomeration at high salt concentrations), and 3) high critical vesiculation concentration (critical concentration of fatty acids required for spontaneous vesicle formation), higher than, for example, the equivalent for phospholipid vesicle formation.

The limited pH stability and salt flocculation can be improved through the covalent cross-linking of neighboring chains of fatty acids, which could have been induced by either UV-C light, temperatures above 50°*C*, simple aging, or any combination of these [34, 35]. A structure of side-by-side overlap of chains of the two layers of the bilayer vesicle, with an overlap of double bonds, provides further stability over an even wider pH range (2-14) [1]. This side-by-side structure enhances the likelihood of cross-linking by all three mechanisms mentioned. Considering the light, temperature, pH, and salt conditions of the Archean ocean surface, cross-linking within the vesicle bilayer of about 18-carbon atom fatty acids would be expected to have occurred.

### 2.4 Optical Properties of fatty acids vesicles

#### 2.4.1 Scattering

When electromagnetic radiation impinges on a particle in-bedded in a medium, the particle’s material polarizes. The induced dipoles oscillate with the incoming radiation and produce a new electromagnetic field, known as the scattered field. The incoming light may also be absorbed by the particles material. The particle thus has both a real and imaginary refractive index. Our fatty acid vesicle is such a particle in-bedded in ocean surface water with sunlight incident from above. As a first approximation, we assume the vesicle to consist of a spherical core made of DNA, RNA, and other fundamental molecules in a water solution, covered with a bi-layer membrane of 18-carbon atom fatty acids (Fig. 2).

**Figure 2.**
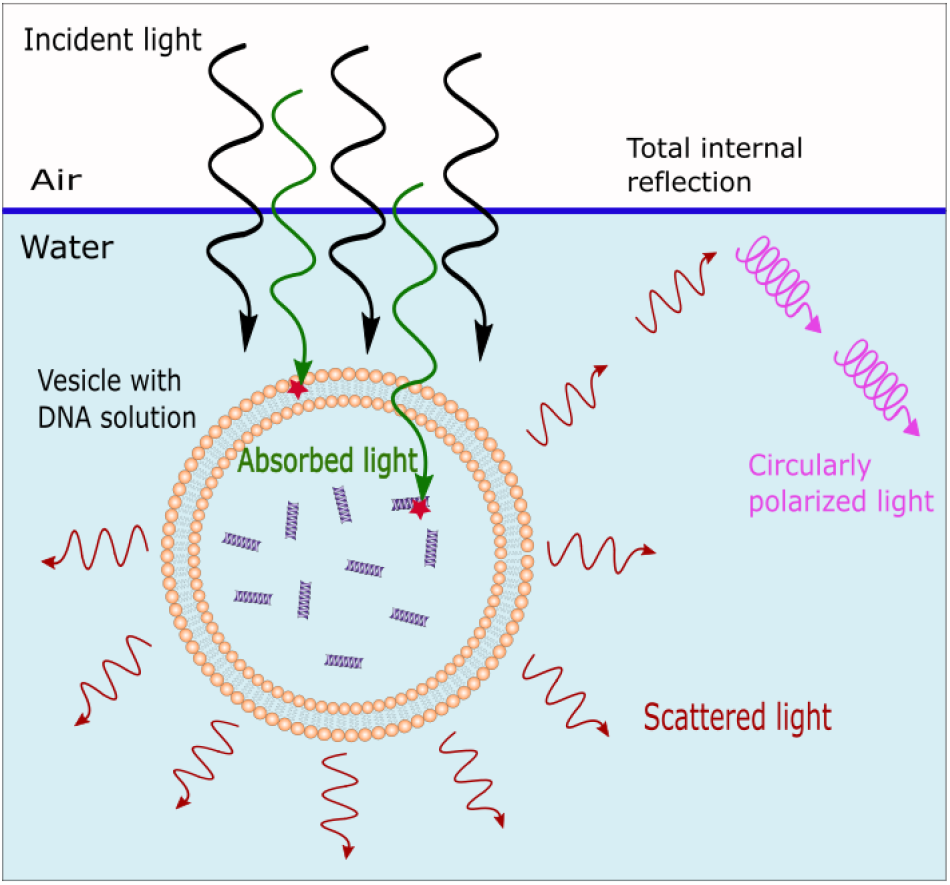
Scattered and absorbed light by a fatty acids vesicle. Light from the sun travels through a thin layer of ocean water and interacts with the vesicle. Some of the light is absorbed by the fatty acids and the DNA, RNA, and other fundamental molecules assumed to have been dissipatively structured within the vesicle. Another part of the light is scattered, mainly in the forward direction. The scattered light is plane polarized and some of it may then be totally internally reflected at the ocean surface, producing circularly polarized light which may have been important in giving rise to the homochirality of life [14].

The efficiencies *Q* of extinction, scattering, and absorption (extinction minus scattering) are defined as the ratio of the observed cross-sections to the geometrical cross-section, which, for a spherical vesicle of radius *a*, is *πa*^2^ [36]. We employed the BHCOAT code [36] to compute the extinction efficiency, *Q_ext_*, the scattering efficiency, *Q_sca_*, the absorption efficiency, *Q_abs_*, and the backscattering efficiency, *Q_back_*, for the vesicle for unpolarized light. The computation assumes Mie theory, valid for vesicles larger than the incident wavelengths.

Input to the BHCOAT code for calculating Mie scattering consisted of the incident light wavelength, the radius of the vesicle core, the thickness of the fatty acid membrane (estimated to be 5 nm for a bilayer of 18-carbon atom fatty acids), the real refractive index of the surrounding water medium, and the complex refractive indexes of the fatty acids and the core solution containing DNA, RNA and other fundamental molecules in water. The real refractive index *n* for water as a function of wavelength was taken from the data of Hale and Querry [37]. The real refractive index of the core solution of DNA, RNA and other fundamental molecules in water depends on the concentration of these (see figure 5 of reference [38]) and was taken to have a range of between 1.01 and 1.02 times that of pure water at 589 nm (*n* = 1.34) giving a range of n between 1.353 and 1.367, which corresponds to the range measured for present day cell nuclei [39]. The refractive index of the fatty acid vesicle wall was taken to be 1.39 at 589 nm [40]. The wavelength dependence of the real refractive index of the core solution and of the fatty acid vesicle wall were taken to be the same as that of pure water multiplied by their corresponding factor, and are plotted in figure 3.

**Figure 3.**
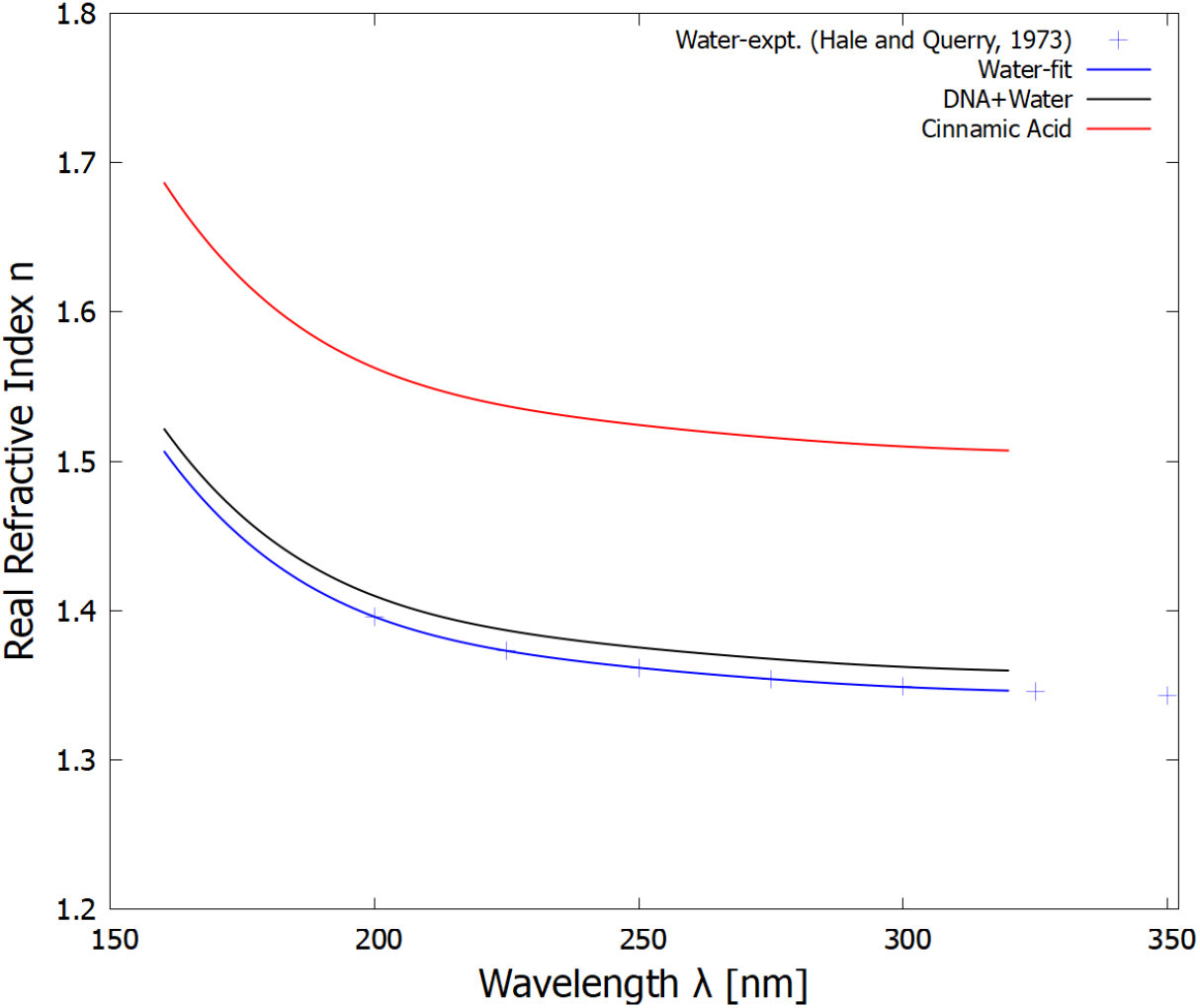
The real part of the refractive indexes for water, the solution of DNA, RNA and other fundamental molecules in water, and the fatty acid bilayer as a function of wavelength. The blue crosses are the experimental data for water obtained from [37]. The refractive index for the fundamental molecules in water was obtained by multiplying that of water by a constant factor of either 1.01 (plotted) or 1.02 (not plotted). The real refractive index of the fatty acid vesicle wall was anchored to a value of 1.39 at 589 nm, corresponding to cinnamic acid [40] (which would be similar to linolenic acid) and given the same wavelength dependence as that of water. Note that because of the small thickness of the vesicle wall (5 nm), the refractive index of the vesicle wall, or its wavelength dependence, does not affect the scattering by any noticeable amount.

The imaginary parts of the complex refractive indexes for water, and for the solution of the fundamental molecules in water, were calculated from the absorption data for water and for DNA in water, respectively, and are plotted in figure 4.

**Figure 4.**
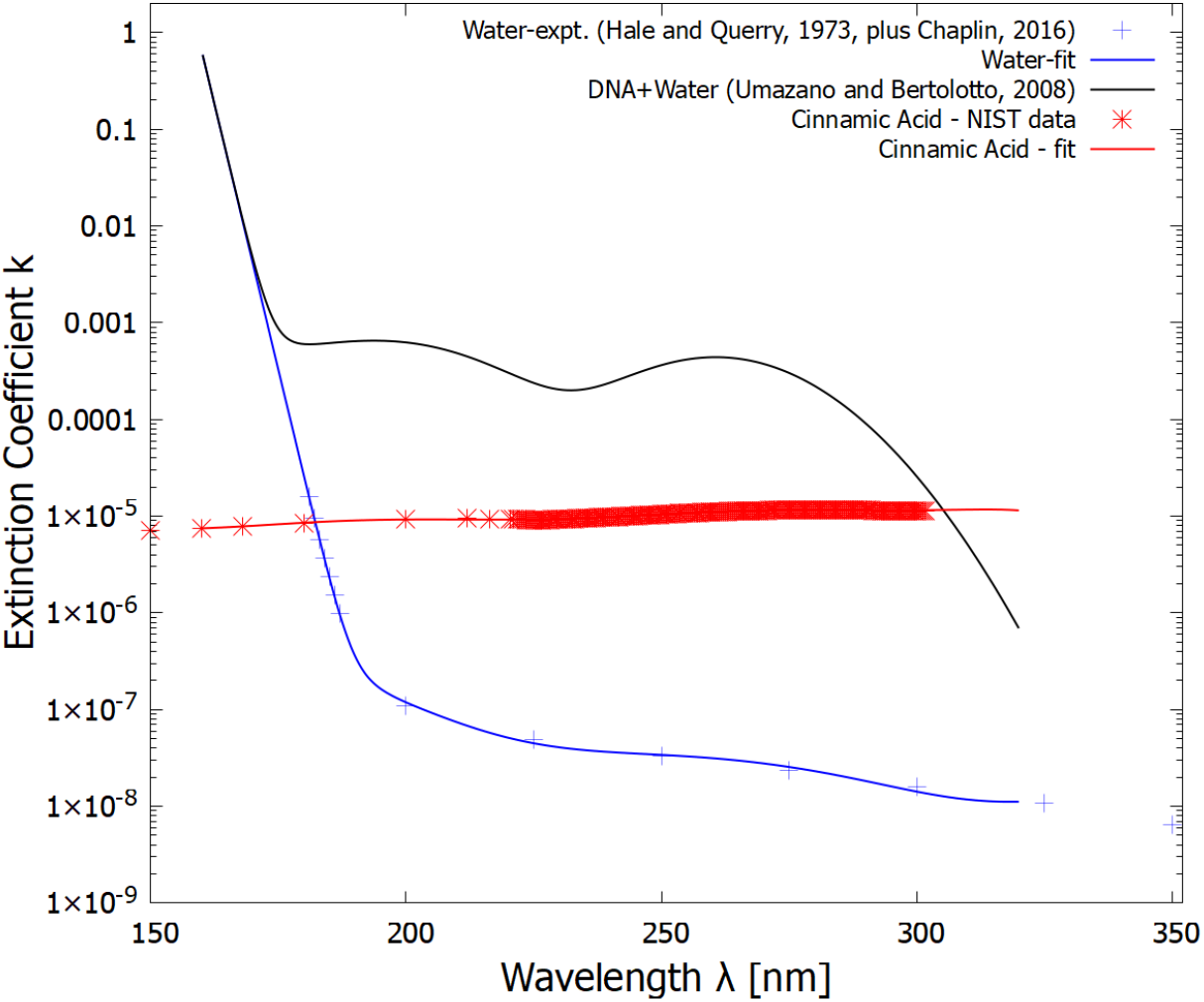
The imaginary part of the refractive indexes for pure water (blue line fit to experimental data of Hale and Querry [37] and Chaplin [41] - crosses), the solution of DNA in water (black line - obtained by multiplying the data from Umazano and Bertolotto [42] by a factor of 10) which represents DNA+RNA+other fundamental molecules in water solution within the vesicle, and the fatty acid bilayer wall (red line - obtained by fitting to data from the NIST handbook for cinnaminic acid, which would be similar to linolenic acid) as a function of wavelength.

The efficiencies *Q_ext_, Q_sca_* and *Q_abs_* (*Q_ext_* – *Q_sca_*) were integrated over the soft and hard UV-C wavelength regions (per unit area presented to the photon beam) plotted as a function of vesicle core radius for the case of the core DNA+fundamental molecule real refractive index of 1.01 times that of pure water in figure 5. The same, but for a core refractive index of 1.02 times that of pure water is given in figure 6. The results show an expected diffraction pattern for small radii. A photo-protective effect against the hard UV-C light is also visible for certain vesicle radii since *Q_sca_* is maximum for the integrated region of 180-210 nm (blue curve, Fig. 5) when the vesicle core radius is 4.4 *μ*m, with another peak at 11.8 *μ*m. These hard UV-C wavelengths have enough energy to dissociate nucleic acids, amino acids, proteins, and other fundamental molecules of life confined within the vesicle. The shielding by Mie scattering thus protects these fundamental molecules from disassociation allowing them to be efficient at absorbing and dissipating in the soft UV-C (245-275 nm) region (Fig. 1). Note, however, that at these small radii, very little of the soft UV-C light is absorbed by the core, most of this light is scattered.

**Figure 5.**
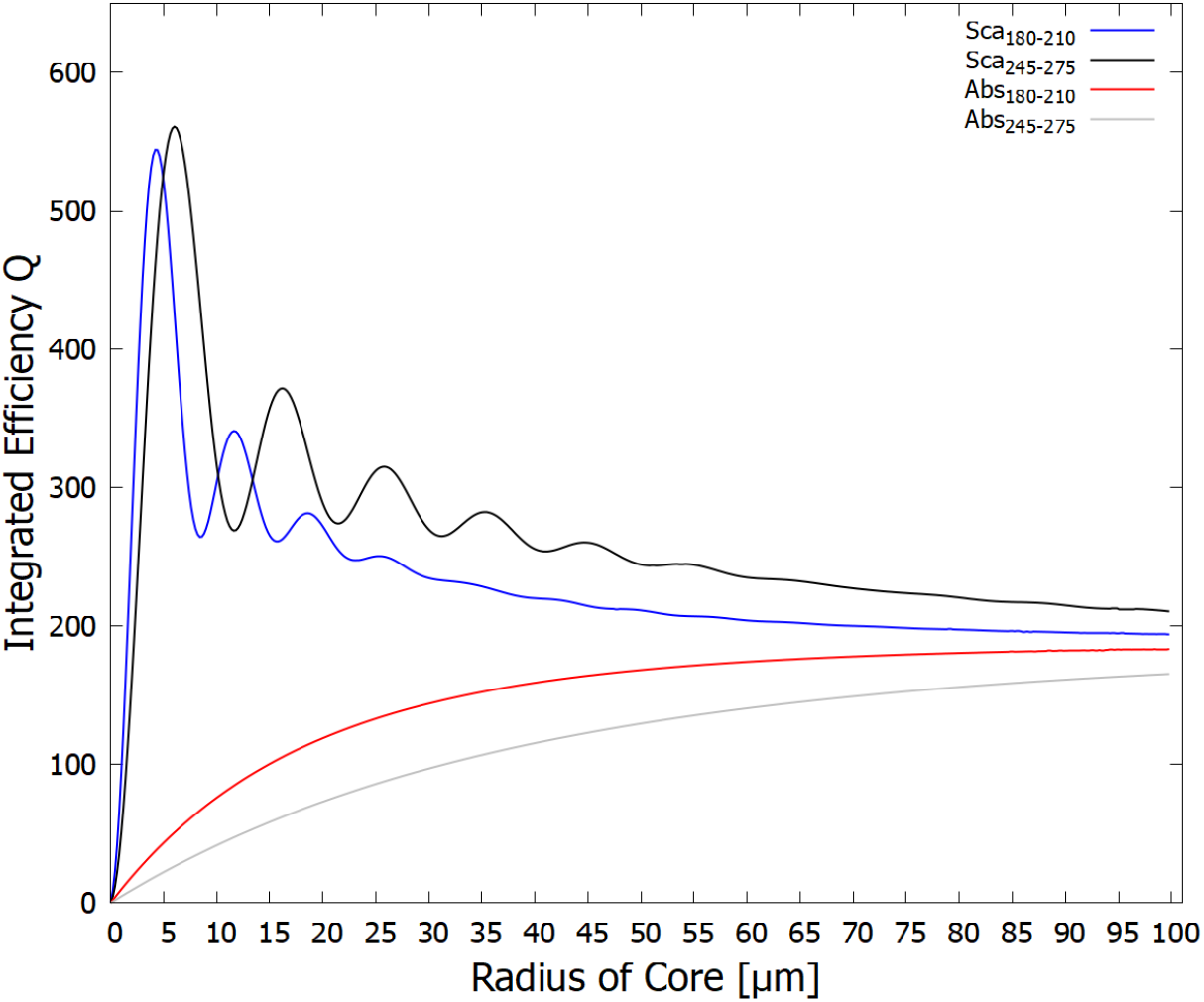
The integrated efficiency of scattering *Q_sca_* and absorption *Q_abs_* per unit area presented to the beam of the fatty acid vesicle for two different wavelength regions (180-210 nm - ionizing/disassociation region) and (245-275 nm - dissipative structuring region) as a function of vesicle core radius (the fatty acid vesicle wall has a thickness of 5 nm). The real refractive index of the core DNA+fundamental molecule solution is taken to be 1.01 times that of pure water (Fig. 3). Note that the greatest scattering (shielding) of the dangerous ionizing radiation (blue line) occurs for the vesicle radius of about 4.4 *μ*m with a smaller peak at 11.8 *μ*m, which would correspond to protocells of diameter 8.8 and 23.6 *μ*m respectively.

**Figure 6.**
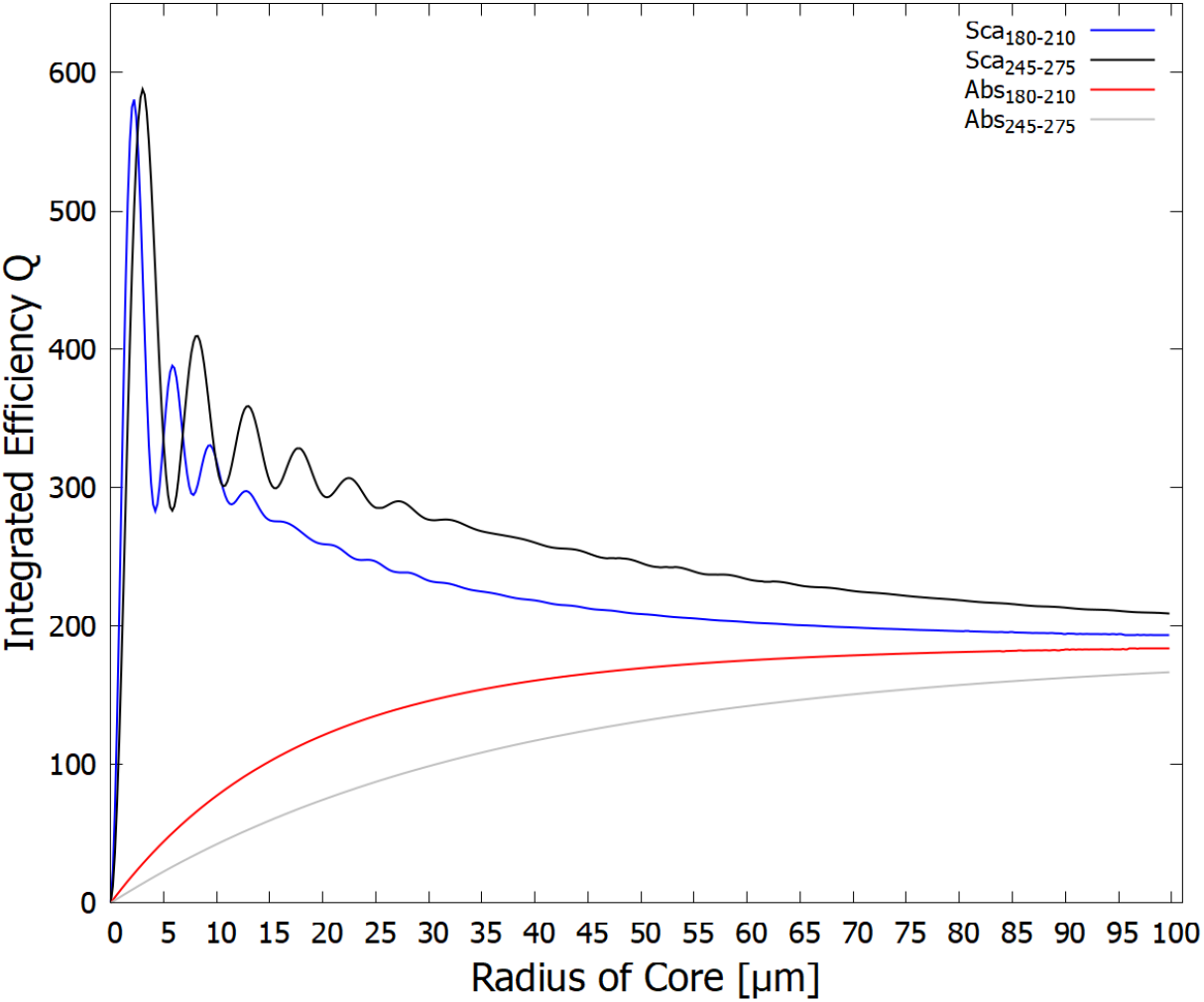
The same as for figure 5 except using a real refractive index for the core DNA+fundamental molecule solution of 1.02 (instead of 1.01) times that of pure water. The greatest scattering (shielding) of the dangerous ionizing radiation (blue line) now occurs for a smaller vesicle radius of about 2.2 *μ*m with a smaller peak at 5.9 *μ*m, corresponding to protocell diameters of 4.4 and 11.8 *μ*m respectively.

In figures 7 through 9, we plot the wavelength dependence of the extinction, scattering, and absorption efficiencies (observed cross-sections compared to the geometrical cross-section of the vesicle, i.e. *πa*^2^) for different-sized vesicles. The fact that the efficiencies can be greater than 1 (and the fact that there is a diffraction-like pattern at small radii, results from the fact that photons are not point particles but really quantum waves interacting with the vesicle edge [36]. Figure 7 corresponds to the vesicle radius (4.4 *μ*m) at which the integrated scattering over the dangerous ionization/disassociation region is maximum (see blue line, Fig. 5). The scattering in the dangerous region of 180 to 210 nm is about 2.8 times greater than that which would be expected classically given the geometrical cross-section. Such vesicle radii would thus be useful, for example, during epochs in which there was lower CO_2_ or little H_2_S (ejected from volcanoes) in the atmosphere to shield the fundamental molecules from this dangerous radiation (see Fig. 1). Note, however, that the absorption (red line) in the soft UV-C region (245-275 nm) is quite small since strong scattering in this spectral region also occurs.

**Figure 7.**
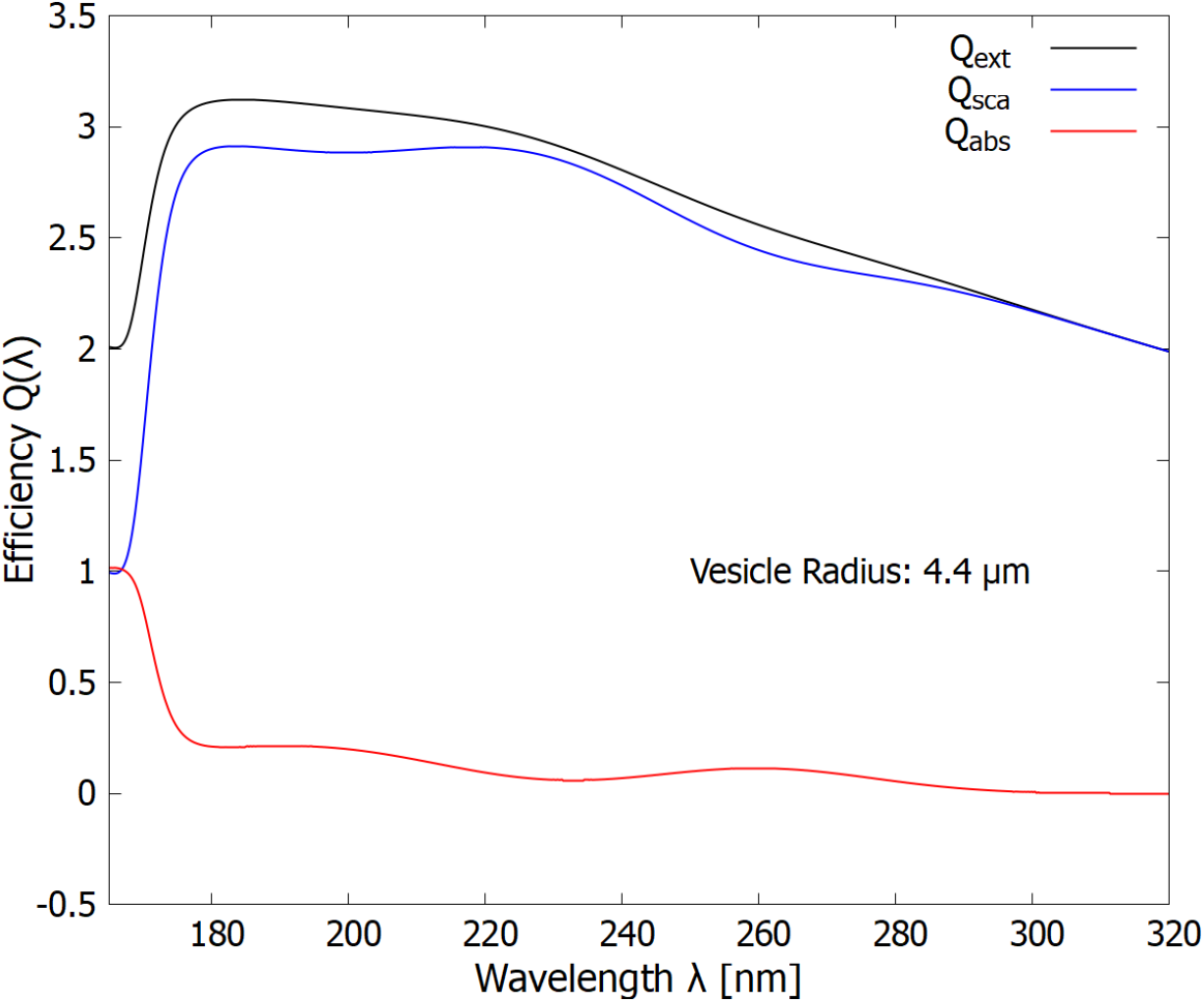
The wavelength dependence of the extinction, scattering, and absorption efficiencies (cross sections compared to the geometrical cross-section of the vesicle) for a vesicle size of 4.4 *μ*m giving maximum protection in the dangerous ionization/disassociation region of 180 to 210 nm (see figure 5).

A second, smaller maximum in the scattering of the dangerous hard UV-C photons is observed at 11.7 *μ*m (Fig. 5). The wavelength dependence of this scattering is given in figure 8. At these greater radii, less, but still significant, scattering occurs in the dangerous hard UV-C region (about 1.75 times that of the geometrical cross-section) but more light is absorbed in the soft UV-C region useful for dissipative structuring.

**Figure 8.**
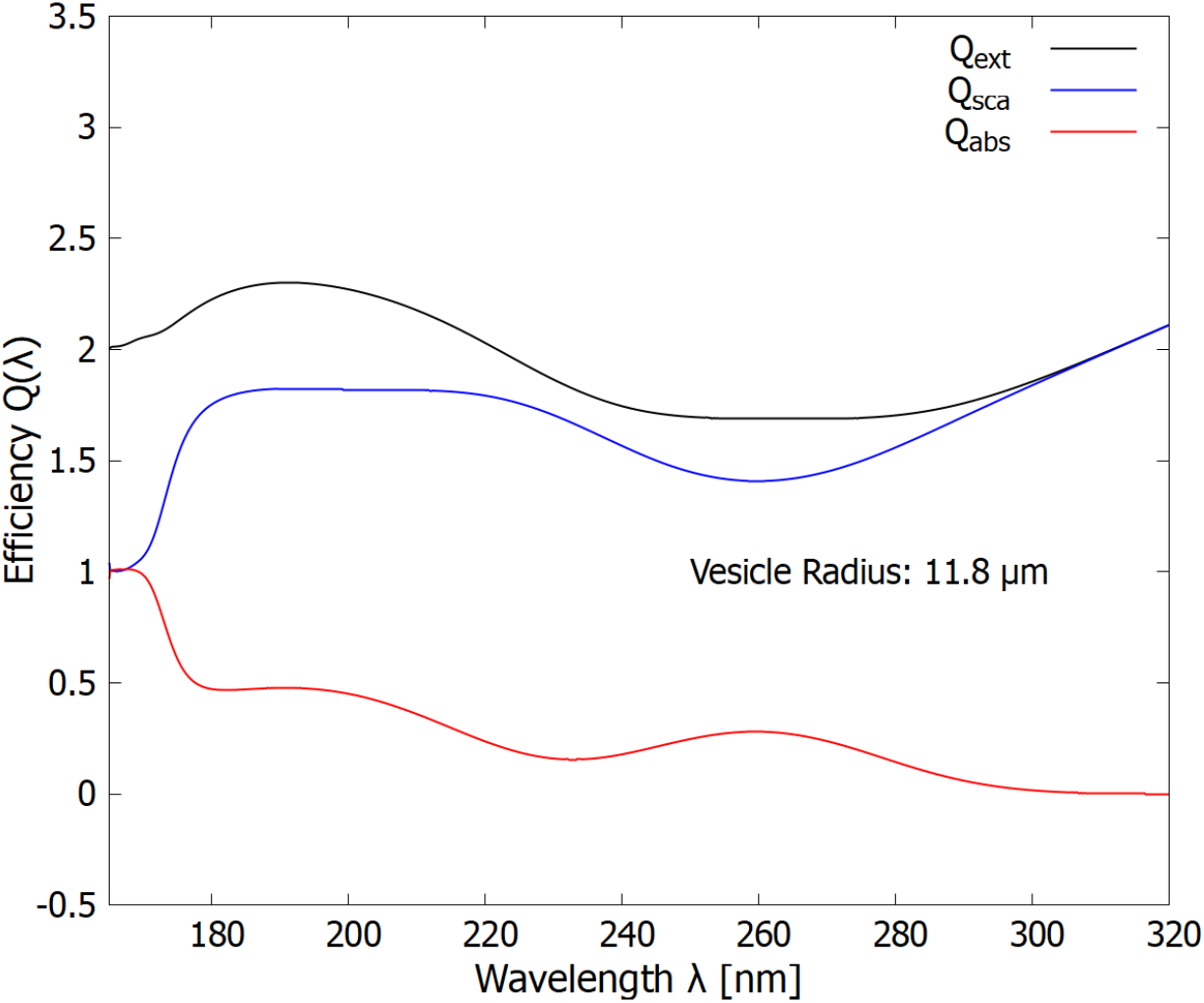
The wavelength dependence of the extinction, scattering, and absorption efficiencies (cross sections compared to the geometrical cross section of the vesicle) for a vesicle radius of 11.8 *μ*m. The scattering (blue line) protection in the ionizing/disassociation region (180-210 nm) decreases somewhat (compare with Fig. 7) while absorption (red line) increases in the dissipative structuring region (245-275 nm).

At the largest radius studied here (100 *μ*m), the scattering and absorption in the hard UV-C region are approximately equal and we find a maximum in the absorption of the soft UV-C photon region. Such vesicle radii would thus be useful (dissipative) when there was a lot of CO_2_ or H_2_S in the atmosphere, or for vesicles at greater depths of the ocean water.

Larger vesicle sizes thus show less protection in the ionization/disassociation region (Fig. 8) but more absorption in the soft UV-C region, while certain smaller sizes show greater protection in the ionization/disassociation region but less absorption in the important dissipative structuring region (Fig. 9).

**Figure 9.**
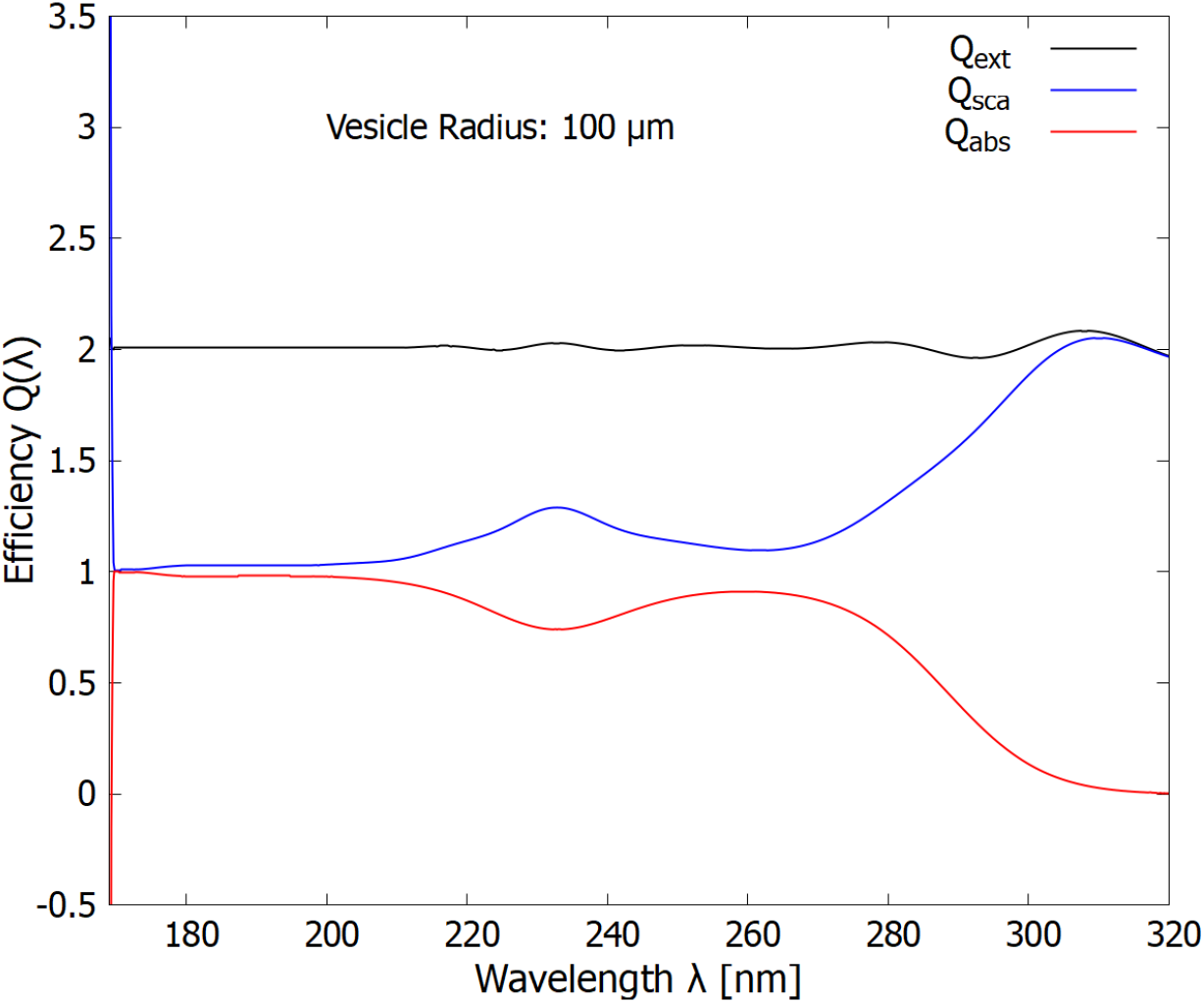
The wavelength dependence of the extinction, scattering, and absorption efficiencies (cross sections compared to the geometrical cross-section of the vesicle) for a vesicle radius of 100 *μ*m (corresponding to the high end of bacteria sizes of today). The Mie scattering (blue line) protection in the ionizing/disassociation region (180 to 210 nm) decreases significantly while absorption in the dissipative structuring region (245-275 nm) increases significantly.

From the above figures and discussion, it is apparent that early life could have tuned the vesicle radius for optimizing the survival and proliferation of the fundamental molecules to the existing light conditions, which probably varied somewhat throughout the Archean. The same could also be acheived by affecting the concentration of fundamental molecules in their interior (thereby changing the core refractive index - compare figures 5 and 6) by, for example, affecting the permeability of the vesicle wall.

#### 2.4.2 Backscattering, Optical Dichroism, and Homochirality

A first scattering of light causes an initially unpolarized beam to become linearly polarized. Total internal reflection of this linearly polarized beam at the ocean surface then results in a component of circular polarization (Fig. 2). Depending on the direction of observation, the beam is either left or right-handed circularly polarized [14]. The surface of the ocean is, in fact, known empirically to be the region on Earth with the greatest circular polarization of light, reaching up to 5% of the available submarine light at the ocean surface for low solar angles to the zenith [43].

We have demonstrated [14] that given the circular dichroism of the nucleic acids, and the measured circular polarized component of light today just beneath the ocean surface, and assuming a similar component during the Archean, and given the existence of ultraviolet and temperature-assisted denaturing of double strand DNA or RNA [16], complete 100% homochirality could have been produced in as little as a few thousand Archean years [14].

In figure 10 we show the integrated backscattering of the soft UV-C photons (245-275 nm) as a function of vesicle radius. Backscattering is greatest at large vesicle radii, as would be expected classically. This backscattered light could then be totally internally reflected from beneath the ocean surface with the sun overhead. At the shallower solar angles, of the morning and afternoon, a much greater amount of forward scattered light could also be totally internally reflected at the surface, giving a much greater component of circular polarization. A considerable amount of light from either backscattering at higher solar angles, or from forward scattering at shallower solar angles, from vesicles at the surface could have contributed to the formation of homochirality through UV-C and temperature assisted denaturing [16] in the nucleic acids within neighboring vesicles.

**Figure 10.**
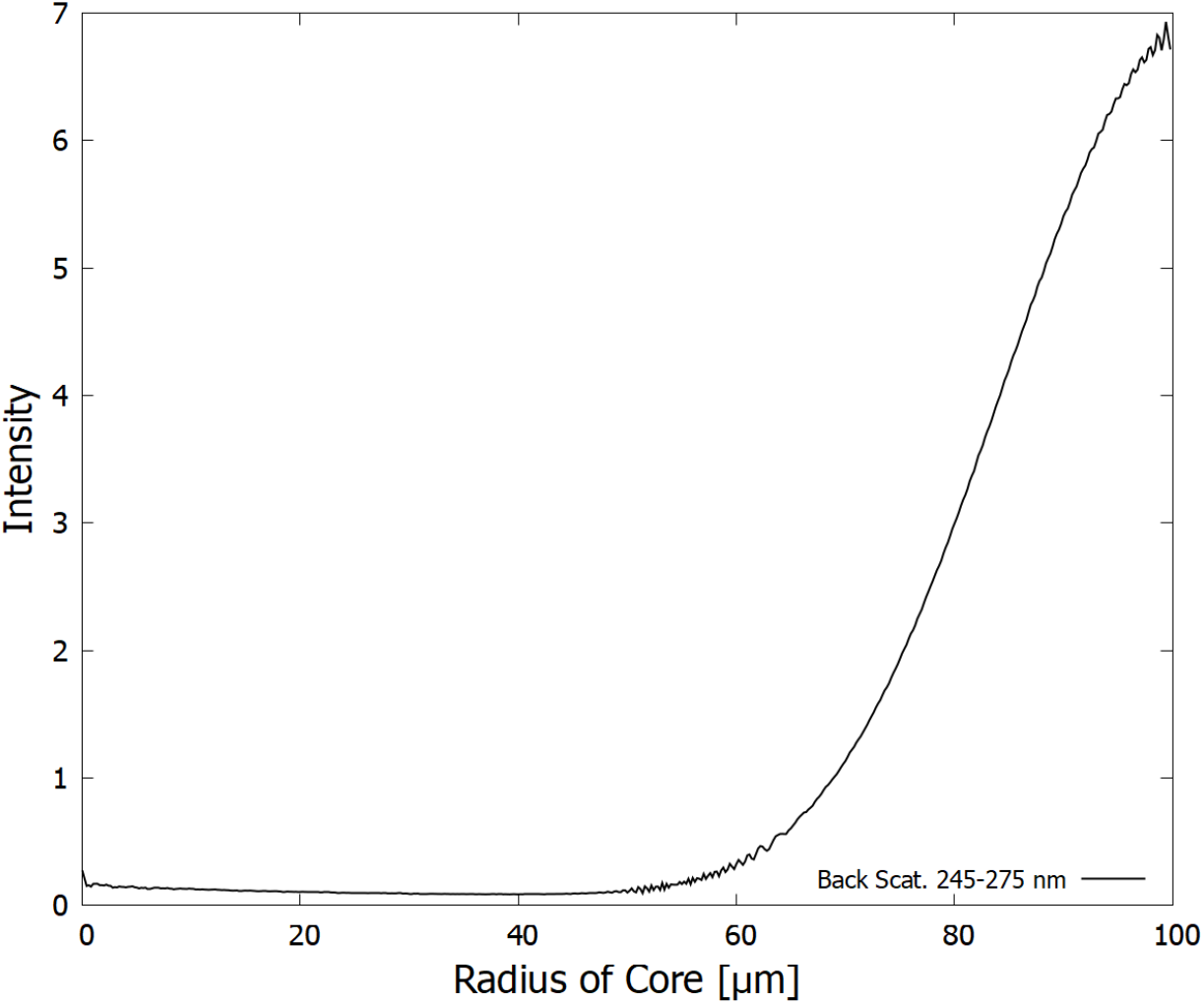
The efficiency of backscattering *Q_back_* into a unit solid angle for the fatty acid vesicle for the wavelength region (245-275 nm - dissipative structuring region) as a function of vesicle core radius (the fatty acid mantel has a thickness of 5 nm and the core solution of fundamental molecules and water has a refractive index of 1.01 times that of pure water). Note that backscattering is an order of magnitude smaller than forward scattering and relatively independent of angle. It increases the larger the vesicle radius. This backscattered light, along with some forward scattered light at low solar angles, if then totally internally reflected at the ocean surface would provide circularly polarized light for inducing the homochirality of the nucleic acids in neighboring vesicles [14, 16].

## 3 Discussion and Conclusions

The thermodynamic dissipation theory for the origin and evolution of life [5, 8, 9, 12, 44] proposes that fatty acids and other fundamental molecules of life were dissipatively structured from simple precursors such as HCN and cyanogen on the ocean surface under the Archean UV-C solar light flux. As for all the fundamental molecules of life, these dissipatively structured fatty acids performed the thermodynamic function of dissipating the UV-C photon potential into heat [10]. Evidence for this includes the fact that conjugated fatty acids absorb strongly this light and have conical intersections allowing the photon-induced excitation energy to be dissipated rapidly into heat. Furthermore, the fatty acids are formed through photochemical routes in which incident photon dissipation rates increase with each step of the photochemical reaction process on route to the final product, which has one or more conical intersections. Such photochemical evolution is the hallmark of molecular dissipative structuring [5, 12].

In this paper we looked, for the first time, at the UV-C optical properties of vesicles made up of a bilayer of fatty acids. In particular, we determined that these properties could have led to some shielding of the damaging hard UV-C photons of the Archean which could have caused ionization and disassociation of the fundamental molecules. According to Mie theory employed here, maximum scattering of the hard UV-C photons by the vesicle occurs at particular radii that depend on the concentration of the DNA and fundamental molecule solution in the core (the real refractive index). These sizes, for concentrations of fundamental molecules similar to that of today’s cells, are consistent with the sizes of fossil bacteria discovered in sediments of the early Archean period [18] when intense UV-C light was arriving at Earth’s surface. Figures 5 and 6 show that the greater the refractive index of the core with respect to that of the surrounding water, the smaller the radius of the vesicle which gives maximum scattering of the damaging hard UV-C photons.

In our analysis, the wavelength dependence of the real refractive index of the DNA+fundamental molecule+water vesicle core solution was taken to be the same as that of water multiplied by a constant factor of either 1.01 or 1.02, giving the range of refractive indexes at 589 nm of present day cells. A better calculation of the refractive index at UV-C wavelengths could be obtained from future experiments or from careful Kramers-Kronig relations given precise wavelength dependent extinction coefficients.

For concentrations of fundamental molecules similar to those found in modern-day bacteria (giving real refractive indexes about 1.01 times that of pure water), maximum hard UV-C scattering occurs at small vesicle diameters (¡ 24 *μ*m), while maximum absorption of soft UV-C photons occurs at diameters greater than 200 *μ*m. The size of the vesicle could thus have been adjusted to give the greatest dissipative structuring with the least amount of molecular disassociation through ionization according to the particular surface UV-C photon spectrum of the epoch (dependent on the pressure of the atmospheric gases CO_2_ and H_2_S which absorb the hard UV-C photons). For example, vesicles of unicellular bacteria have been found in Archean deposits with sizes of less than 24 *μ*m at ~ 3.43 Ga, while with dimensions of up to 289 *μ*m at ~ 3.2 Ga [18].

Backscattering of light with the sun at high angles, or forward scattering at low solar angles, of the soft UV-C photons incident on the vesicle, could have been totally internally reflected at the ocean surface, resulting in a component of circularly polarized light which could have induced homochirality in the nucleic acids of neighboring vesicles [14, 16].

The results obtained here imply that fatty acid vesicles could indeed have shielded UV ionizing radiation from interfering with the molecular dissipative structuring of the fundamental molecules during the Archean. Such structuring may thus also be viable on planets of any stars emitting significant UV-C light, even though the planet may not contain sufficient atmospheric CO_2_ or hydrogen sulfide during a given epoch to filter out all the high energy ionizing and disassociating photons. This would make the origin of life a more robust process, and open up the possibility that carbon based life as we know it may exist on a much greater number of planets.

## Abbreviations

**The following abbreviations are used in this manuscript:**

CO_2_: carbon dioxide
DNA: deoxyribonucleic acid
HCN: hydrogen cyanide
H_2_S: hydrogen sulfide
RNA: ribonucleic acid
UV-A: light in the region 360-400 nm
UV-B: light in the region 285-360 nm
UV-C: light in the region 100-285 nm (only the region 180-285 nm is relevant here since shorter wavelengths are well shielded by atmospheric CO_2_)
UVTAR: Ultraviolet and Temperature Assisted Replication

## Acknowledgments

This research was funded by DGAPA-UNAM, grant number IN104920. I.L. is also grateful for a national postgraduate scholarship from CONACyT. This document is based on the [arXiv / bioRxiv Template](https://www.overleaf.com/latex/templates/arxiv-slash-biorxiv-template/phncddwqtxpc) by Philipp Schlegel under the license [CC BY 4.0](https://creativecommons.org/licenses/by/4.0/).

